# Global atlas tree of natural proteins based on sorted composition vectors

**DOI:** 10.1101/866103

**Authors:** Pu Tian

## Abstract

Sequence comparison is the cornerstone of bioinformatics and is traditionally realized by alignment. Unfortunately, exponential computational complexity renders rigorous multiple sequence alignment (MSA) intractable. Approximate algorithms and heuristics provide acceptable performance for relatively small number of sequences but engender prohibitive computational cost and unbounded accumulation of error for massive sequence sets. Alignment free algorithms achieved linear computational cost for sequence pair comparison but the challenge for multiple sequence comparison (MSC) remains. Meanwhile, various number of parameters and procedures need to be empirically adjusted for different MSC tasks with their complex interactions and impact not well understood. Therefore, development of efficient and nonparametric global sequence comparison method is essential for explosive sequencing data. It is shown here that sorted composition vector (SCV), which is based on a physical perspective on sequence composition constraint, is a feasible non-parametric encoding scheme for global protein sequence comparison and classification with linear computational complexity, and provides a global atlas tree for natural protein sequences. This finding renders massive sequence comparison and classification, which is infeasible on supercomputers, routine on a workstation. SCV sets an example of one-way encoding that might revolutionize recognition and classification tasks in general.

## Introduction

The ability to compare and classify sequences is essential for modern biology. Its importance is partially reflected by the fact that BLAST/X^1,2^ and CLUSTALW(X)^3,4^ ranked 12, 14, 10 and 28 respectively among most cited scientific papers^5^. Sequence comparison is expected to become more critical as single cell sequencing technologies mature. Traditionally, alignment is the way to compare sequences and has a rich history of development^1–4,6–8^. Unfortunately, rigorous MSA is computationally intractable. Approximate MSA algorithms^7,9^ work well for small number of sequences (e.g. less than 50,000). Alignment free algorithms^10–13^, while reduce computational cost of a single sequence pair comparison to linear complexity with respect to chain length, do not resolve the daunting task of MSC. Besides the prohibitive computational cost of MSA/MSC, extent of error propagation for present approximate algorithms is unknown for massive sequence sets. The need for empirical parameters (e.g. scoring and gap parameters for alignment and word length for alignment free algorithms), MSA/C heuristics and complex interactions among these factors are additional source of uncertainties for present methodologies. Impact of these issues are likely to become more severe for massive sequence sets that are expected to be generated routinely. Therefore, efficient, reliable and non-parametric MSC method is highly desired. A global atlas for natural proteins, if successfully constructed and enriched with our routine sequence alignments/comparisons, would enable us marching toward a full understanding of the protein fold space and evolutional history. It is the intent of this work to start a first step toward these directions.

Present sequence comparison algorithms are universally applicable for any sequences on arbitrary alphabets (with necessary tuning of parameters/heuristics), such general ability inevitably reduces its efficiency for special type of sequences, with biological sequences being an outstanding example. Natural protein sequences are subjected to strong cellular environmental, synthesis, structural, functional, dynamical and degradation constraints, and are presumably to be confined in a low dimensional continuous (if we believed in evolution) manifold within vast possible sequence space (approximately 20^301^ as estimated by Uniref50 sequence set). One corollary is that highly efficient one-way encoding schemes (that discard significant manifold boundary information but keep intra-manifold differences) are likely to exist for biological sequences. Therefore it is possible to first project natural sequences to a dramatically compressed space where different individuals within the unknown manifold can still be distinguished to simplify the MSA/MSC challenge. The sorted composition vector (SCV) encoding is proposed with this thought in mind, and is found to be useful in realizing non-parametric qualitative global protein sequence comparison and classification for arbitrarily large sequence set (that can be stored in memory).

### Molecular constraints and sorted composition vector encoding

With given environment, chemical composition is usually the most important property of a molecule in determining its behavior. From a coarse-grained perspective, all natural proteins swim in similar biological media (despite its complexity and diversity). Intra-molecular environment of each amino acid (AA) is mainly determined by composition of the belonging chain. One intuitive way is to designate the most populous AA type as the most important composition, and the second most populous AA type to be the second most important composition, etc. With this idea, Natural protein sequences may be easily mapped into sorted composition vectors (SCVs) with lengths equal to or shorter than 20. This immediately partition the whole protein sequence space into a tree with 6613313319248080000 nodes (see Figure 1). Natural proteins may range from small peptides to over 30000-AA long chains and with the average length being approximately 301 for the uniref50 data set (see Methods). Thus a potential sequence space of approximately 20^301^ is reduced by SCVs to less than 10^19^. The specific intra-molecular environment hopefully specifies behavior of each comprising AA, hence that of the chain. It is no doubt that identical sequences share the same SCV. Our analysis suggest that intra-molecular constraints specified by SCV, together with stringent extra-molecular constraints experienced by natural proteins, dictate that (in overwhelming majority cases) only very similar sequences may share the same SCV, which thus provides an ultra efficient encoding scheme. The resulting SCV tree (see figure 1) generates a global atlas for natural proteins, and realizes qualitative MSC for massive sequence sets with no adjustable parameters. Construction of such an atlas tree takes a one-pass reading of all available sequences, thus have an effective time and space computational complexity of *NL* for *N* chains with length *L* (or *L*_1_ + *L*_2_ + … + *L*_*N*_ for *N* chains with various lengths). When compared with present MSA/C/ algorithms with typical *N*^3^ time complexity scaling and *N*^2^ space complexity scaling for consistency based methods^7^, it is about *N*^2^ (or more) times faster, that is 10^12^ faster for a sequence set of size one million, and has much less memory requirement.

**Figure 1.**
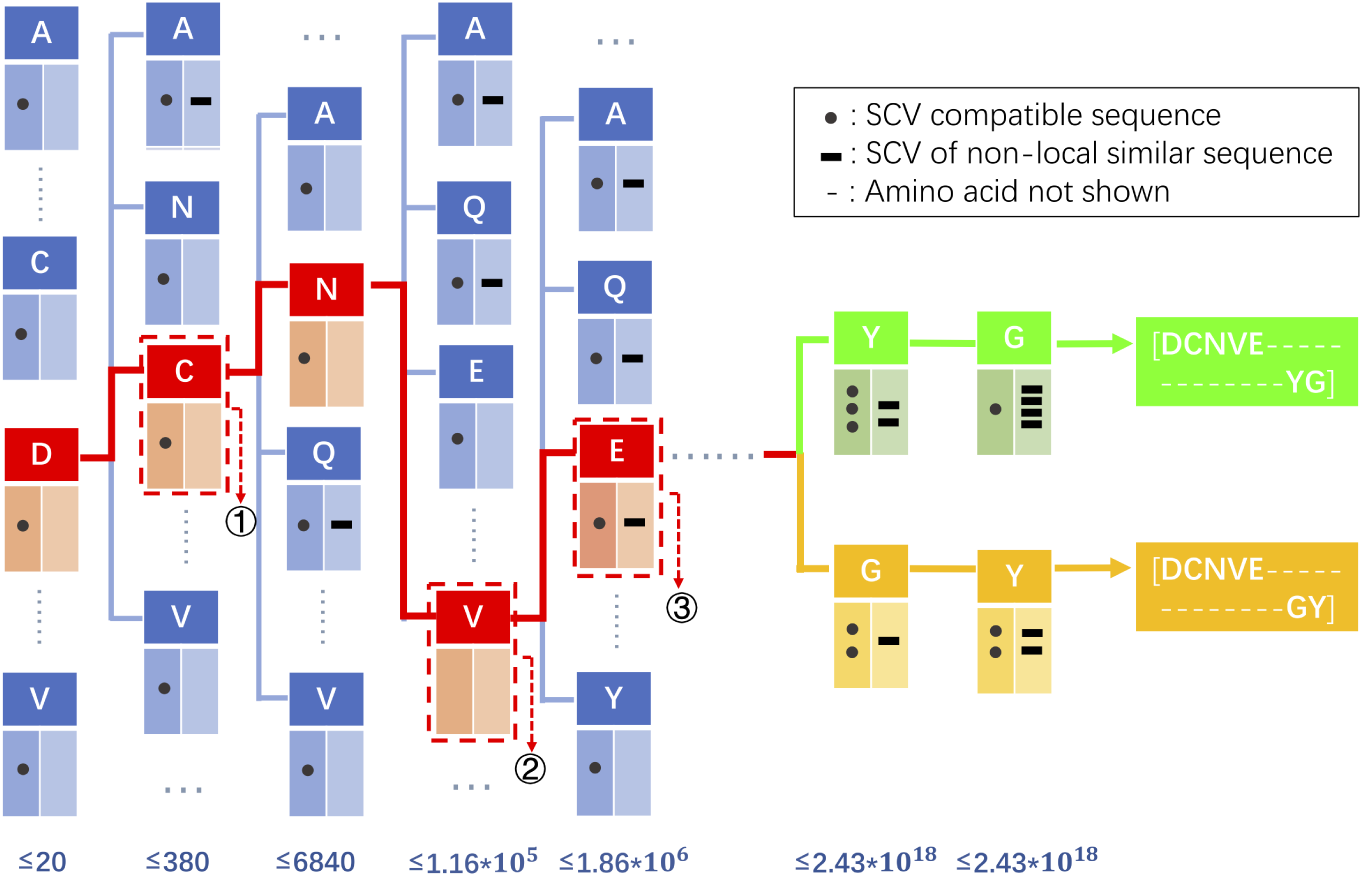
Schematic representation of a protein SCV tree with 20 layers (shown from left to right) and a maximum possible of 6613313319248080000 nodes (*N*_*nodes*_ = 20!/(20 −*n*)! for the *n*th layer), only partial tree is presented with only one child branch (except the last two layers) shown for each layer in the figure with omitted part denoted by dots. Each node has three boxes, the title node is essential and is denoted by a letter in corresponding SCV, each letter of which specifies its path in the tree; sequence box save the list of sequences (denoted by circles) encoding the SCV that ends with the title node letter and with other letters specified by ancestor nodes; distant neighbor box saves non-local SCV nodes (denoted by rectangles) that containing similar sequences as those in the sequence box. There are three types of nodes (denoted by circled numbers), 1) nodes with a sequence list, 2) path nodes with no sequences and 4) nodes with both sequences and SCV list of distant similar sequences. The first level of this tree has 20 nodes, each of which represent homo-polypeptides of arbitrary length with the same weight vector which is simply [1.0]. The second level correspond to protein chain comprising two types of amino acid with the majority being specified by its parental node. For example ′*AAL*′ and ′*AAAALL*′ share the same node (*AL*) in the SCV tree with the same weight vector [⅔, ⅓]. Thus SCV and its accompanying weight vector provide very little constraints for homo-polypeptides and di-hetero-polypeptides. Fortunately, the overwhelming majority (more than 99.63% in uniref50 and 99.85% in uniref100) of natural proteins have more than 13 different types of comprising AA (see Table 1). Two SCV with 20 types of AA [*DCNVE* −−−−−−−−−−−−−*YG*] and [*DCNVE* −−−−−−−−−−−−−*GY*] are partially shown in the figure with hyphens represent omitted AA.

### Distribution of similar protein sequences in SCV tree

When more than one protein sequences encode the same SCV, corresponding nodes are termed multiples. SCV trees for two different data sets (uniref50, uniref100, see Methods) are constructed. Number of tree paths, sequences, filled nodes and multiples for various SCV lengths are listed in table 1. Most filled nodes have only sequence, and overwhelming majority of rare multiples have only two chains (*N*_*seq*_ − *N*_*Fnode*_ is only slightly larger than 2 × *N*_*MTP*_ in table 1). Sequences in the same multiple have very high sequence identity (Figures 2 and 3). Removal of identical (or similar) chains in a massive sequence set is trivial in SCV tree as only comparison within rare multiples (or limited number of neighborhood and annotated distant nodes) is necessary. Sequences in uniref50 are supposed to have pairwise sequence identity lower than 50% as expected by CD-HIT algorithm^14^, which was utilized to remove similar sequences from uniref100 to construct uniref50. The fact that some very similar sequences remain in uniref50 is due to unbound propagation of local sequence comparison error. No present algorithm can guarantee full removal of highly similar sequences in a massive dataset without a full pairwise comparison, which is about *N/*2 (*N* is the number of sequences in the set) times more expensive than SCV tree construction and does not provide qualitative global relative relationship as revealed by SCV tree.

**Table 1.** Number of paths (*N*_*Path*_), sequences *N*_*Seq*_, filled nodes (*N*_*FNode*_) and multiple nodes (*N*_*MTP*_) in each layer of the SCV tree built from uniref50 and uniref100.

| UniRef50 |  |  |  | UniRef100 |  |  |  |
| --- | --- | --- | --- | --- | --- | --- | --- |
| $N_{Path}$ | $N_{Seq}$ | $N_{FNode}$ | $N_{MTP}$ | $N_{Path}$ | $N_{Seq}$ | $N_{FNode}$ | $N_{MTP}$ |
| 20 | 0 | 0 | 0 | 20 | 0 | 0 | 0 |
| 380 | 16 | 15 | 1 | 380 | 82 | 54 | 14 |
| 6536 | 45 | 39 | 5 | 6605 | 335 | 155 | 34 |
| 84887 | 158 | 120 | 17 | 90847 | 1018 | 469 | 85 |
| 663290 | 304 | 285 | 13 | 802077 | 1645 | 1107 | 167 |
| 2989062 | 633 | 613 | 14 | 4245091 | 2588 | 2063 | 226 |
| 8561826 | 1077 | 1063 | 10 | 14552648 | 3996 | 3357 | 241 |
| 16934103 | 1945 | 1932 | 12 | 35277439 | 6011 | 5149 | 273 |
| 25056216 | 3533 | 3519 | 10 | 64303785 | 9083 | 8235 | 340 |
| 30584132 | 6684 | 6666 | 18 | 94150371 | 16075 | 14234 | 620 |
| 33628295 | 13325 | 13305 | 17 | 118919774 | 28955 | 26652 | 1066 |
| 35180597 | 29827 | 29792 | 29 | 137545396 | 62752 | 58445 | 2129 |
| 35872847 | 72530 | 72496 | 32 | 150989634 | 148004 | 139982 | 4378 |
| 36078849 | 179625 | 179581 | 41 | 159816809 | 363626 | 350165 | 8967 |
| 35993895 | 422712 | 422650 | 57 | 164760431 | 900242 | 869273 | 20948 |
| 35600912 | 916062 | 915977 | 82 | 166810238 | 2129682 | 2058765 | 48733 |
| 34694127 | 1843625 | 1843517 | 97 | 166351670 | 4883674 | 4701479 | 125789 |
| 32853550 | 3618282 | 3618128 | 148 | 162480418 | 12191326 | 11456020 | 433134 |
| 29236678 | 7845291 | 7844932 | 354 | 151452229 | 38463028 | 34613993 | 2014086 |
| 21392008 | 21395233 | 21392008 | 3070 | 116974828 | 135988894 | 116974828 | 8945458 |

**Figure 2.**
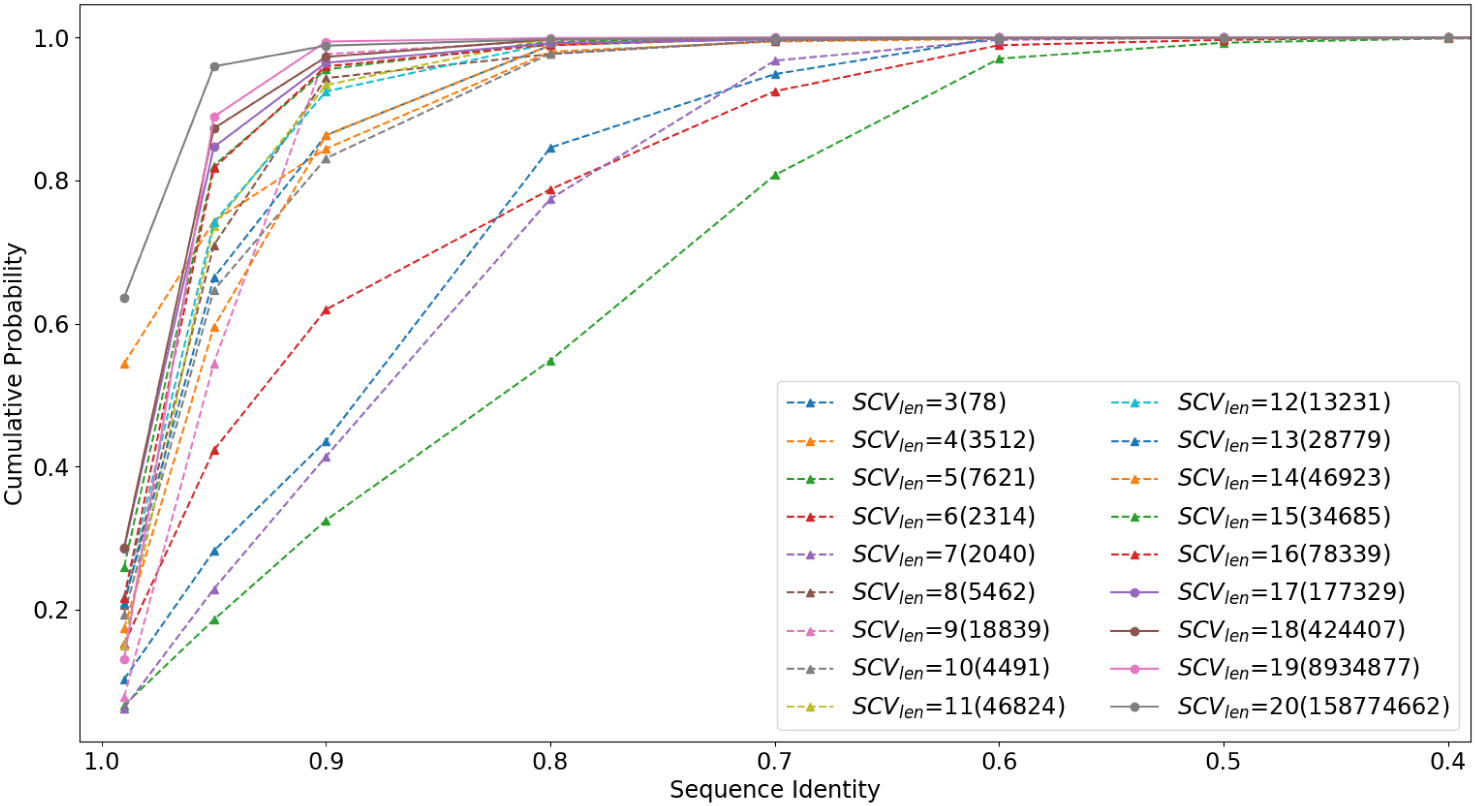
Cumulative probability of within-multiple sequence pair identity for SCV lengths from 3 to 20. Number of calculated sequence pairs is shown in parenthesis. Data points of X-axis are [0.99, 0.95, 0.9, 0.8, 0.7, 0.6, 0.5, 0.4]. The value at 0.99 is the percentage of within-multiple sequence pairs that have sequence identity higher than 99%, etc. For SCV lengths larger than 7, the overwhelming majority within-multiple sequences have pair identity higher than 0.8. For SCV lengths larger than 16, the overwhelming majority within-multiple sequences have pair identity higher than 0.9. The results are based on Uniref100 set.

**Figure 3.**
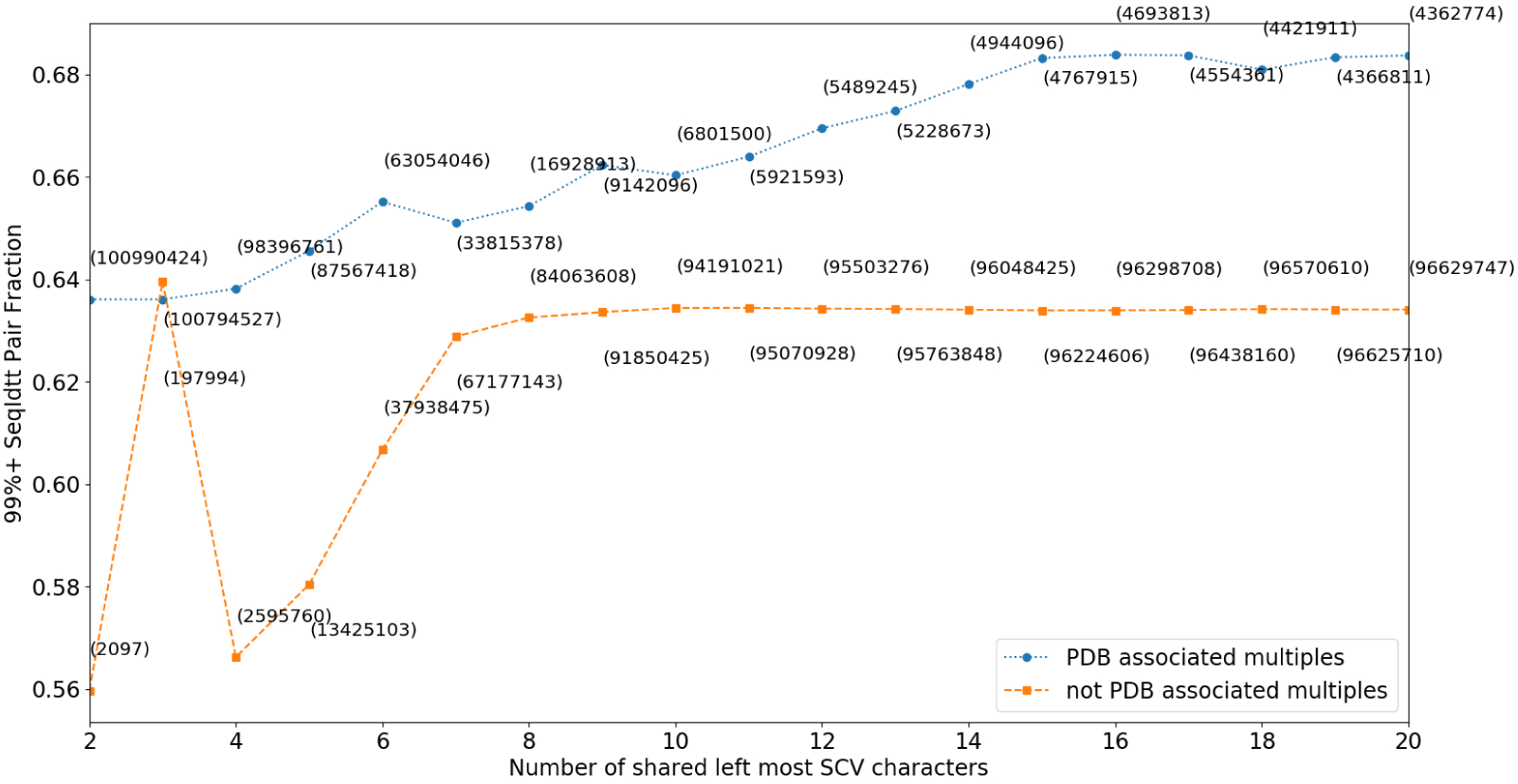
Percentage of within-multiple pair sequence identity higher than 99% for all 20-letter SCV sharing (circles) and not-sharing (squares) various number of left most letters with sequences in PDB database.

**Figure 4.**
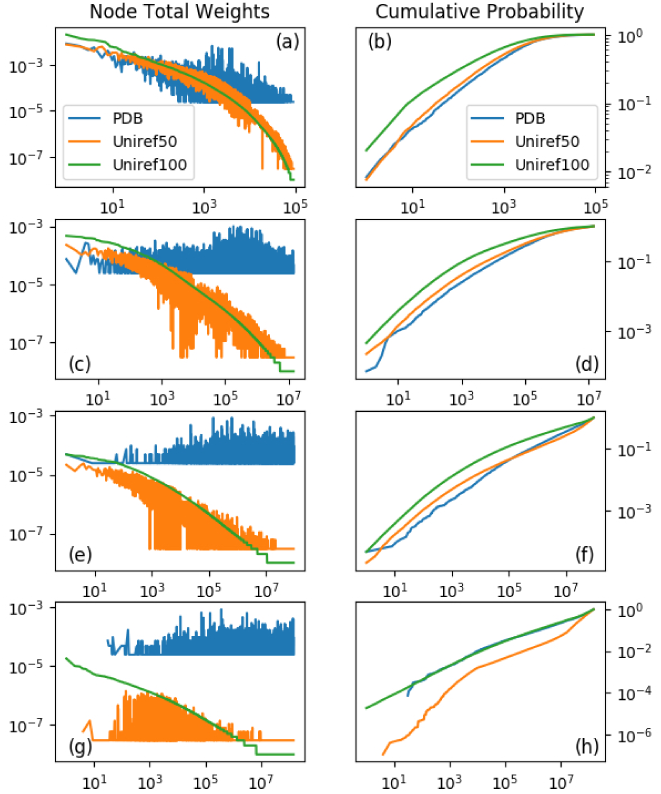
Distribution (a,c,e,g) and cumulative probability (b,d,f,h) of sorted total node weight defined by Uniref100 sequence set. Total sequence number of a node is the sum of number of sequences fall in a node and all of its descendent nodes. Total node weight is normalized total sequence number for all nodes at the same level of SCV tree. a) and b) are for the 3 − *rd* level nodes; c) and d) are for the 6*th* level nodes; e) and f) are for the 9*th* level nodes; g) and h) are for the 12*th* level nodes

As shown in Fig. 2, the extent by intramolecular constraints of given SCVs increase rapidly with their lengths such that qualitatively sequences in multiples correspond to longer SCV exhibit significantly higher sequence identity (the reason this relationship is not strictly observed is likely due to lack of statistics), and the relative number of multiples (*N*_*MTP*_*/N*_*FNode*_) are smaller for nodes of longer SCVs when SCV length is smaller than 13. It is important to note that while overwhelming majority of chains have SCV longer than 13, SCV tree path branching does not increase significantly after the 13th layer. Within a SCV multiple node, weight vectors provide more quantitative information beyond SCV itself, one may choose to investigate the full sequences since multiples are rare and number of distinct sequences within multiples are small. Besides those reside in the same multiple, sequences reside in closely distanced nodes in SCV tree (e.g. sibling nodes with common close ancestor, parent and child nodes, grand-parent and grand-child nodes) also have quite high sequence similarity. The distance between nodes in SCV tree is a qualitative nonlinear measure for difference of corresponding sequences (with exceptions explained and resolved below). As the branching of SCV tree mainly occurs within the top 13 layers, All SCVs sharing the first 13 letters are likely to be quite similar, and quantitative relationship need to be analyzed further. An additional limitation of the SCV tree is that the extent of differences among all nodes sharing the same parent is not specified, which physically corresponding to addition of a new AA type in the intra-molecular environment specified by the parent node. The tree structure (parent nodes and paths) explicitly specifies conditions of adding in new AA type, thus provides a composition specific log table to record accurate description of quantitative sequence similarity. Full quantitation of SCV tree for natural proteins implicates accomplishing full comparison of natural protein sequence space, and needs to be improved for years to come as statistics accumulates with more available sequences.

For a given sequence, apart from sequences in closely neighboring nodes of SCV tree, there is possibility that some similar sequences may locate far away in the SCV tree. For example, “*ALSGKDEFNQPY*” and “*LASGKDEFNQPY*” are very similar SCVs but locate 24 steps away in the SCV tree. Existence of similar sequences in distant SCV nodes caused by such partial reversal situations (*AL* vs. *LA*) may be annotated for all chains in SCV tree (Fig. 1). For each chain, only a small number of different SCV reversals are possible and therefore their treatment takes linear computational cost.

### Reduction of SCV tree by equal weight amino acid type

When two or more different amino acid type have the same composition weight, their rank become meaningless and could be artificially different depending upon the sorting algorithm and their order of appearance as a chain is read from the amino-terminus to the carboxyl-terminus. To remove such uncertainty, rank of equal weight AA need to be specified. In this work, more populous AAs (as revealed by statistics from the uniref50 sequence set) are placed in front of less populous ones in case of equal weight. Another direct impact is that SCV space (number of nodes in SCV tree) is reduced. The extent of SCV space reduction is estimated on the uniref50 and uniref100 data set respectively and the results are shown in Table 2. Better SCV space reduction estimation may be achieved as more sequences become available.

**Table 2.** Number of protein class nodes for specified length of SCVs. These numbers are calculated from by *N*_*nodes*_ = 20!/(20 − *n*)!

| $SCV_{length}$ | $N_{node}$ | $N_{node\_r\_uni50}$ | $N_{nodes\_r\_uni100}$ |
| --- | --- | --- | --- |
| 1 | 20 | 20.0 | 20.0 |
| 2 | 380 | 380.0 | 354.254 |
| 3 | 6840 | 6232.0 | 6298.92 |
| 4 | 116280 | 1.10269e+5 | 1.03486e+5 |
| 5 | 1860480 | 1.36552e+6 | 1.31214e+6 |
| 6 | 27907200 | 1.53539e+7 | 1.56203e+7 |
| 7 | 390700800 | 1.65610e+8 | 1.64728e+8 |
| 8 | 5079110400 | 1.69212e+9 | 1.63279e+9 |
| 9 | 60949324800 | 1.54313e+10 | 1.43448e+10 |
| 10 | 670442572800 | 1.16505e+11 | 1.02145e+11 |
| 11 | 6704425728000 | 7.13125e+11 | 6.29664e+11 |
| 12 | 60339831552000 | 3.37604e+12 | 3.09156e+12 |
| 13 | 482718652416000 | 1.35510e+13 | 1.36603e+13 |
| 14 | 3379030566912000 | 5.08334e+13 | 5.38978e+13 |
| 15 | 20274183401472000 | 1.82261e+14 | 2.08672e+14 |
| 16 | 101370917007360000 | 6.93837e+14 | 8.84451e+14 |
| 17 | 405483668029440000 | 3.12220e+15 | 4.59311e+15 |
| 18 | 1216451004088320000 | 1.62390e+16 | 2.83181e+16 |
| 19 | 2432902008176640000 | 8.88886e+16 | 1.42621e+17 |
| 20 | 2432902008176640000 | 2.43479e+17 | 2.78874e+17 |

### Physical constraints variation for different SCVs

As is shown in Fig.**??** that more compositions (nodes correspond to longer SCVs) on average correspond to stronger constraints as reflected by larger fraction of within-multiple sequence pairs have higher than 99% sequence identity. Correspondingly, it is intuitive to speculate that SCV multiples with significant in-node sequence differences correspond to relatively weaker composition/physical constraints and allow more different ways of primary sequence permutation variation. As a corollary, protein sequences fall in such SCV nodes are likely to be more flexible and consequently tend to be more difficult to crystallize. Therefore, it is likely that statistically, multiples in SCV tree that share a significant fraction of SCV with any chains in protein data bank (PDB) should on average have lower probability of observing dissimilar chains. This is indeed the case as shown in Fig.**??**, where sequences in multiples sharing various number (from 2 to 20) of first SCV letters with any PDB chains have higher probability of observing higher than 99% sequence identity for in-node pairwise alignment than sequences in those multiples that do not. Similarly, when sequence clusters sharing first *K*(*K* = 3, 6, 9, 12) letters of their SCV code for uniref100 data set are sorted and plotted in Fig.**??**a), c), e) and f). It is shown that the largest (least restrictive) clusters have no PDB chains. Cumulative probability (Fig.**??**b), d) and f)) show that PDB chains are underrepresented for the least restrictive (or most populous) partial SCV codes except when *K* = 12 or larger *K* (not shown). This is consistent with the fact that tree branching does not increase significantly after the 12*th* layer.

## Discussion

While we constructed a global atlas for natural proteins based on SCV in an efficient, robust and nonparametric way. The issue of evaluating similarity of constituting alphabets members in normal sequence alignment persist locally. Since SCV tree by itself does not specify extent of similarity for sequences located in sibling nodes. The fundamental contribution of this work is to limit such uncertainties to local, thus effectively limiting the global propagation of error. Such local similarity is apparently dependent upon compositions specifying corresponding locality in the SCV tree, quantitative characterization of these locality necessitate construction of scoring schemes of some form. However, the great thing is that SCV tree provides an effective global log table, which does not dependent on adjustable parameters, for us to accumulate detailed quantitative analysis of specific local similarities.

The starting point of SCV encoding is utilization of complex inter- and intra-molecular restraints that shape the unknown manifold of natural protein sequences in vast sequence space. When sufficient number of sequences is available, the rank for number of total sequences (in a node and all of its descendent nodes) for the 20 top level gives a qualitative description of relative constraint power of the corresponding AA type, with the AA defining node of the lowest rank (least number of total sequences) has the strongest restraints. Similarly, at each level, relative restraint power of different AA type under the given environment specified by ancestor nodes is reflected by local rank of total sequences. SCV is a specific case of one-way encoding based on unknown manifold.

While the exact encoding scheme utilized here is specific to protein, the general idea might well be applicable to other applications. For example, extension of SCV encoding to nucleotide sequences is likely to be straight forward. SCV vector of maximum lengths (4, 16, 64, 256, 1024, 4096 …) may be used. Specifically, with 5′ to 3′ order, construction of 4 element SCV vector is just a counting of *A, G,C, T*, which is simply the GC/AT content weights and has been widely utilized in genome sequence analysis. However, extension to 16 element SCV can be easily encoded by counting of 16 different dinucleotides, extension to 64 element SCV uses tri-nucleotide segment, etc. Encoding of a nucleic acid sequence into SCVs of various maximum length can be achieved with a single pass of reading. This immediately open the door for massive genome sequence comparison.

Apart from analysis of protein sequences as demonstrated in this work and analysis of nucleic acid (NA) sequences as proposed above, SCV is fundamentally a general one-way encoding scheme that can potentially be applied in encoding of two or higher dimensional data such as pixel images. For example, a two dimensional pixel matrix (or higher dimensional tensor quantity) may be first flattened in a wide variety of mapping order, with value of each pixel discretized as the specific alphabet just as AA. Further generalization of SCV to text, audio and video recognition and classification is possible. Algorithm development and application of SCV in these scenarios will be explored in near future.

Different from traditional compression algorithms that realize two-way encoding, SCV is a one-way encoding scheme with significant irreversible information loss. Essentially, in our specific case of natural protein sequence analysis, it keeps significant differences among in-manifold protein sequences but discards large body of important information on boundaries of the concerned manifold. Apparently, successful inverse mapping from SCV to corresponding original sequences implicates understanding of the lost boundary information for the corresponding manifold, and is a very challenging task. However, as long as a one-way encoding scheme effectively keep in-manifold differences, it is likely to be useful for comparison and classification of interested objects. The utility of one-way encoding is not limited to natural protein sequences (e.g. NA sequences, linearized graphics, text, audio and video objects proposed above). It is likely to be applicable for any low dimensional manifold in high dimensional space as long as proper encoding scheme is constructed. Essentially almost all high dimensional problems in reality might be mapped into a much lower dimensional manifold. Therefore, searching for reliable one-way encoding might be a new direction for robust and efficient objects comparison and classification, which is the central task for many present artificial intelligence (AI) applications. This study is to my best knowledge the first effective proposal and utilization of such one-way encoding. Mathematically, the proposed SCV encoding is an efficient non-linear dimension reduction scheme with linear computational time and space complexity. This is in contrast to widely utilized principle component analysis (PCA), which engenders cubical computational cost to realize linear dimension reduction. The common advantage for SCA and PCA is that both are non-parametric. Extension of SCV in additional (other than discussed above) high dimensional sparse matrix problems in physics, chemistry, machine learning and many other fields is not necessarily straight forward, it might well be inspirational. It is believed by the author that there should be more one-way encoding schemes beyond SCV, and hope that more variety of efficient one-way encoding being developed to target different applications.

## Conclusions

Based on the thought that biological sequences are effectively limited within a low dimensional continuous manifold in a potentially vast high dimensional sequence space, SCV is proposed and demonstrated to be an efficient and reliable encoding scheme that effectively accomplish non-parametric and qualitative global multiple sequence comparison with linear time and space computational complexity. This is in stark contrast to the cubical or higher order time complexity and square space complexity of present available algorithms. SCV provides a physical perspective on intra-molecular constraints that is lacking in all present sequence alignment/comparison algorithms. The atlas tree based on SCV provides an effective log table for future quantitative and global understanding of natural protein universe. Additionally, SCV is an example of one-way encoding that can potentially be applied on other sequential data (e.g. nucleic acid sequences) or higher dimensional data with proper specified flattening procedures (e.g. flattening of 2D or 3D pixel data). Success of SCV suggests that search of effective one-way encoding might be a productive new direction for object comparison and classification in general.

## Methods

We downloaded uniref50.fasta.gz and uniref100.fasta.gz from the uniprot database on Sep/01/2019 and Sep/04/2019. After excluding sequences shorter than 40 and sequences that containing any non-natural AAs, two sequence sets have 36350907 and 195201016 sequences respectively. PDB sequences was downloaded from PDB database. For proteins that have many slightly different sequences due to mutation, only the first one in the original file is selected, similarly, all sequences shorter than 40, have non-natural AAs, and are not included in Uniref100 are removed with 43538 sequences in the final set. After constructing the SCVs for accompanying weight vectors, protein chain IDs are organized into nested python dictionaries. Sequence identity calculation is performed with the Biopython’s Align.Pairwisealigner module.

## Notes and references

[1] S. F. Altschul, W. Gish, W. Miller, E. W. Myers and D. J. Lipman, Journal of Molecular Biology, 1990, 215, 403–410.

[2] S. Altschul, T. Madden, A. Schaffer, J. Zhang, Z. Zhang, W. Miller and D. Lipman, FASEB Journal, 1998, 12, 3389–3402.

[3] J. D. Thompson, D. G. Higgins and T. J. Gibson, Nucleic Acids Research, 1994, 22, 4673–4680.

[4] J. D. Thompson, T. J. Gibson, F. Plewniak, F. Jeanmougin and D. G. Higgins, Nucleic Acids Research, 1997, 25, 4876–4882.

[5] R. Van Noorden, B. Maher and R. Nuzzo, Nature, 2014, 514, 550–553.

[6] S. R. Eddy, Current Opinion in Structural Biology, 1996, 6, 361–365.

[7] M. Chatzou, C. Magis, J. M. Chang, C. Kemena, G. Bussotti, I. Erb and C. Notredame, Briefings in Bioinformatics, 2016, 17, 1009–1023.

[8] H. Ashkenazy, I. Sela, E. Levy Karin, G. Landan and T. Pupko, Systematic Biology, 2019, 68, 117–130.

[9] S. Baichoo and C. A. Ouzounis, BioSystems, 2017, 156-157, 72–85.

[10] I. Schwende and T. D. Pham, Briefings in Bioinformatics, 2014, 15, 354–368.

[11] C. A. Leimeister, M. Boden, S. Horwege, S. Lindner and B. Morgenstern, Bioinformatics, 2014, 30, 1991–1999.

[12] S. Sarmashghi, K. Bohmann, P. M. Gilbert, V. Bafna and S. Mirarab, Genome Biology, 2019, 20, 1–20.

[13] A. Zielezinski, H. Z. Girgis, G. Bernard, C. A. Leimeister, K. Tang, T. Dencker, A. K. Lau, S. Röhling, J. J. Choi, M. S. Waterman, M. Comin, S. H. Kim, S. Vinga, J. S. Almeida, C. X. Chan, B. T. James, F. Sun, B. Morgenstern and W. M. Karlowski, Genome Biology, 2019, 20, 1–18.

[14] L. Fu, B. Niu, Z. Zhu, S. Wu and W. Li, Bioinformatics, 2012, 28, 3150–3152.

